# Designing Effective small interfering RNA for Post-Transcriptional Silencing of Human GREM1: A Comprehensive Bioinformatics Approach

**DOI:** 10.1101/2020.01.23.917559

**Authors:** Asiedu Ebenezer

## Abstract

Human gremlin-1 is a physiologically versatile signaling molecule that has been associated with several human diseases including cancer. The ability of gremlin-1 to induce fibrosis in organs and transduce angiogenesis makes it a target for cancer therapy. RNAi-based therapy has proven to be very efficient and specific in tumor growth inhibition. The efficacy and specificity of siRNA-mediated gene silencing depends on the designing approaches. Here, empirical guidelines for siRNA design and comprehensive target site availability analysis were used to select effective siRNA from a plethora of potential candidates designed using several computation algorithms. Then, the selected siRNA candidates were subjected to stringent similarity searches in order to obtain siRNA candidates with reduced off-target effects (high specificity). The best candidates were compared to experimentally successful gremlin-1 siRNAs in order to predict the silencing potency of the selected siRNAs. siRNA-6 (sense strand: 5’-CCAAGAAAUUCACUACCAU-3’), siRNA-7 (sense strand: 5’-CCAUGAUGGUCACACUCAA-3’) and siRNA-47 (sense strand: 5’-GGCCCAGCACAAUGACUCA-3’) were predicted to be highly effective siRNA candidates for gremlin-1 silencing. These siRNAs can be considered for RNAi-based therapy because off-target effects are predicted to be minimal.

## INTRODUCTION

GREM1, located on human chromosome 15q13→q15, is the gene that codes for gremlin-1. Gremlin-1 is a member of the DAN (differential screening-selected gene aberrative in neuroblastoma) family of BMP antagonists. This family of proteins are highly conserved biomolecules with a cysteine-knot domain and are very active signaling elements during embryonic development [1]. Gremlin-1 is a 184-amino acid protein which plays crucial roles during vertebrate limb patterning [2], cell differentiation [3], organogenesis [4] and angiogenesis [5]. The physiological versatility of gremlin-1 explains why the protein is linked to several human pathologies [6] and cancer progressions [7–9]. Gremlin-1 induces fibrosis in many organs which results in progression of several pathologies including diabetic nephropathy and pulmonary hypertension [10]. Moreover, gremlin-1 mediates thrombo-inflammation in some cardiomyopathies [11]. Over the years, considerable evidence depicting gremlin-1 as an oncogenic and pro-angiogenic factor has been reported. The overexpression of gremlin-1 in tumor cells as well as their neighboring stromal cells has been observed in several carcinomas including those of the breast [12] and colon [13]. Apart from modulating BMP signaling, gremlin-1 can be involved in a BMP-independent signaling via vascular endothelial growth factor receptor-2 (VEGFR-2) to promote angiogenesis [14]. Promoting angiogenesis subsequently enhances the rate of tumor growth and progression. Gremlin-1 regulates the transition of epithelial cells to mesenchymal cells, thus contributing to the progression of cancers [15].

RNA interference (RNAi) is an effective therapeutic strategy to silence the expression of target genes in many remediation processes including cancer therapy. When the target gene is a matured mRNA, the interference mechanism is described as post-transcriptional gene silencing [16]. RNAi is an important biological process in hosts during gene regulation and cell defense. The mechanism of RNAi is mediated by endogenous or exogenous double stranded RNA (dsRNA) together with other complex proteins. The dsRNA is processed into small interference RNA (siRNA) duplexes of about 19-23 bases with 2-3 nucleotide overhangs [17]. The processing of dsRNA to siRNAs is done by a complex protein, Dicer, together with double stranded RNA binding proteins [18]. The dicer mediated processing is followed by an integration of the siRNAs to RNA-induced silencing complex (RISC) by RISC-loading complex (RLC). RISC has an ATP-dependent RNA helicase domain that separates the two strands of the siRNA. The strand whose 5’ end has the lower free energy of binding, the guide strand, is bound by the RISC whereas its complementary strand, the passenger strand, is cleaved and removed. The active siRNA-RISC (si-RISC) complex recognizes its target mRNA. An RNase III component of RISC known as Argonaute-2 cleaves the mRNA opposite the bound guide strand, thereby preventing its translation.

Despite the challenges, siRNA-based therapeutics have seen developments in the clinical setting [19]. Endogenous RNAi machinery processes exogenous siRNA to silence target genes. Major issues in siRNA-based therapy have to do with delivery, efficacy, off-target effects, toxicity and immunostimulatory effects [19]. Careful selection and design of potential therapeutic siRNA can alleviate some of the major challenges especially off-target effects. The success of siRNA knockdown depends on several factors that must be considered during siRNA design. The intrinsic characteristics of the siRNA is of prime importance to knockdown efficiency since minor changes in siRNA sequence could affect the functionality [20]. In view of this, properties such as nucleotide content, sequence length and duplex thermodynamics should be considered during siRNA design. The target binding site is also a subject of evaluation in terms of its location and accessibility [16].

Several guidelines for designing effective siRNA to target mammalian genes have been reported [16, 21]. Online software tools available for siRNA design integrate some of the proposed guidelines in their algorithms to ensure reliable predictions. In this study, the guidelines proposed by Ui-Tei [22], Reynolds [23] and Amarzguioui [24] are used to select effective siRNAs from a number of potential candidates designed by multiple computational algorithms. A comprehensive analysis of target site availabilities for the selected candidates are performed. The siRNA designed here are meant to target human gremlin-1 mRNA, hence, efforts are made to ensure reduced off-target effects. The best candidates selected in this study are compared with experimentally successful siRNAs to determine the gene silencing potency of the selected siRNAs.

## METHOD

### Data collection

Human gremlin-1 primary transcript (NM_013372.7) and protein sequence (ID: O60565) were retrieved from the NCBI nucleotide database (https://www.ncbi.nlm.nih.gov/nuccore) and UniProtKB (https://www.uniprot.org/uniprot/) respectively. The mature mRNA that codes for the 184-amino acid gremlin-1 was analyzed using the ExPASy translate tool (https://web.expasy.org/translate/). Fasta file of the human gremlin-1 primary transcript (NM_013372.7) was the input data. Reading on both forward and reverse strands were allowed and the output format set to include nucleotide sequences. The precise nucleotide sequences (open reading frame) responsible for coding human gremlin-1 was used as the target for subsequent siRNA design.

### In-Silico design of siRNA

Here, three designing algorithms and two commercial design software tools were used to design candidate siRNA to target human gremlin-1. The tools are: OligoWalk (http://rna.urmc.rochester.edu/RNAstructureWeb/), RNAxs (http://rna.tbi.univie.ac.at/cgi-bin/RNAxs/RNAxs.cgi), i-score Designer (https://www.med.nagoya-u.ac.jp/neurogenetics/i_Score/i_score.html) and the commercially available Invitrogen BLOCK-iT™ RNAi Designer (https://rnaidesigner.thermofisher.com/rnaiexpress/) and Invivogen siRNA Wizard (https://www.invivogen.com/sirnawizard/). The OligoWalk algorithm predicts siRNA candidates considering the thermodynamics of RNA hybridization [25]. The server predicts siRNA candidates based on the free energy changes upon binding of an siRNA to target mRNA and intrinsic nucleotide sequence characteristics. RNAxs predicts siRNA candidates that satisfy proposed rules for siRNA design and also have target sites that are highly accessible [26]. For the i-score algorithm, a linear regression model examines each siRNA nucleotide, then generate a scoring argument to calculate the inhibitory score (i-score). The scoring argument informs which nucleotides are favored at which positions considering proposed nucleotide-based rules for siRNA design [27].

### Selection of effective siRNA using proposed guidelines for siRNA design

The guidelines proposed by Ui-Tei, Reynolds and Amarzguioui were used to manually select the best siRNA candidates using a scoring method (Table 1). The scoring method was designed to focus on nucleotide sequence composition only. Guidelines addressing duplex thermodynamics and siRNA folding were excluded from the scoring method, however, were addressed in subsequent analysis. siRNA candidates that scored at least 75% (23 out of 30) were selected for further analysis.

**Table 1:** Scoring method used for grading siRNA candidates based on preferred nucleotide sequence features.

| Ui tei et al., 2004 |  | Amarzguioui et al., 2004 |  | Reynolds et al., 2004 |  |
| --- | --- | --- | --- | --- | --- |
| rule | score | rule | score | rule | score |
| The 10th and 19th nucleotide of the sense strand should be A or U | 2 | Absence of U at 1st position and G at 19th position of the sense strand | 2 | U at position 10 in the sense strand | 2 |
| AU rich at the 5' end (7 bases) of the antisense strand | 2 | A nucleotide at 6th position in the sense strand | 2 | At least 3 A/U bases in the region of 15–19th bases | 2 |
| The sense strand should have more than 3 A/U bases in the region between the 13th and 19th bases | 2 | GC content should be moderate 32–58% | 2 | Absence of internal repeats | 2 |
| The 1st base of the sense strand should be G/C | 2 | A/U at position on 19th base of the sense strand | 2 | The 3rd and 19th base of the sense strand should be A | 2 |
|  |  | Presence of C nucleotide at 16th position of sense strand | 2 | The 13th base of the sense strand should not be G | 2 |
|  |  | Presence of U at 13th position of sense strand | 2 | The stable hairpin-like secondary structure should be avoided | 2 |

### Target site accessibility prediction

The accessibility of the siRNA target sites was analyzed using the Sfold server (http://sfold.wadsworth.org/cgi-bin/index.pl). The Sirna module was particularly used for this activity. The Sirna module is for rational design of siRNA considering proposed rules for siRNA design, duplex thermodynamics and target site accessibility [28]. Given a target mRNA as input, the algorithm evaluates accessibility of the target site using parameters such as probability profiles and loop specific profiles. Internal stability profiles of siRNA for particular target positions were also analyzed.

### Similarity search for siRNA candidates

The standard nucleotide blast (blastn) of the NCBI (https://blast.ncbi.nlm.nih.gov/Blast.cgi) was used to search for sequences that share homology with the selected siRNA candidates. The human Genomic plus Transcript (Human G + T) database was used for all the similarity searches. The blast algorithm was optimized to search for somewhat similar sequences (blastn) for all siRNA sequences and mega-blast for mRNA sequence. Expect threshold was set to 1000 and scoring parameters set to default.

## RESULTS

### Design and Selection of effective siRNA candidates

Gremlin-1 primary transcript containing 14575 nucleotides was analyzed with the ExPASy translation tool. The open reading frame (ORF) that codes for the human gremlin-1 was predicted to be in 5’ 3’ Frame 1 (Supplementary Fig. 1). The ORF was observed to be continuous stretch of nucleotides starting from position 160 to 757 of the primary transcript (Supplementary Fig. 2). Several in-silico tools were used to design siRNA candidates that target gremlin-1 mRNA, precisely the ORF sequence. Sixty-six potential siRNAs were designed to silence gremlin-1 (Supplementary Table 1). The most effective and best siRNA candidates were selected using empirical guidelines proposed by Ui-Tei et al., Reynolds et al., and Amarzguioui et al. The scores for all the designed siRNAs are shown in Supplementary Table 2. The researchers reported a strong correlation between siRNA silencing efficacy and nucleotide sequence characteristics. As shown in Table 1, each guideline describes features that are required for effective siRNA-mediated gene silencing, including mammalian genes [22]. High scoring siRNA candidates therefore exhibit enough nucleotide-based features required for effective gene silencing. As shown in Table 2, four siRNA candidates scored at least 75%. The ideal length of siRNA for effective gene silencing is still a subject of controversy in the design of siRNA. The dicer-mediated processing of dsRNAs results in short sequences (19-23) of siRNA with 2 nucleotide overhangs. Longer siRNA sequences (25-27) have also been reported to be functional because the dicer will process such sequences [29]. The GC content of an siRNA duplex is a parameter that can define the gene silencing efficacy of siRNAs. A moderate occurrence of GC in the siRNA (≈32%-≈58%) is preferential [24]. GC content of the high scoring siRNAs ranged from 34% to 58% (Table 2). A high GC% delays unwinding of siRNA duplex by nuclear helicases whereas a low GC% slows hybridization of siRNA to target mRNA [24]. Thermodynamics properties of duplex ends also define the silencing efficacy of siRNA in cells [30]. The internal stability of the high scoring siRNA candidates was evaluated with the Srna module available at Sfold and the results are shown in Figure 2.

**Table 2:**
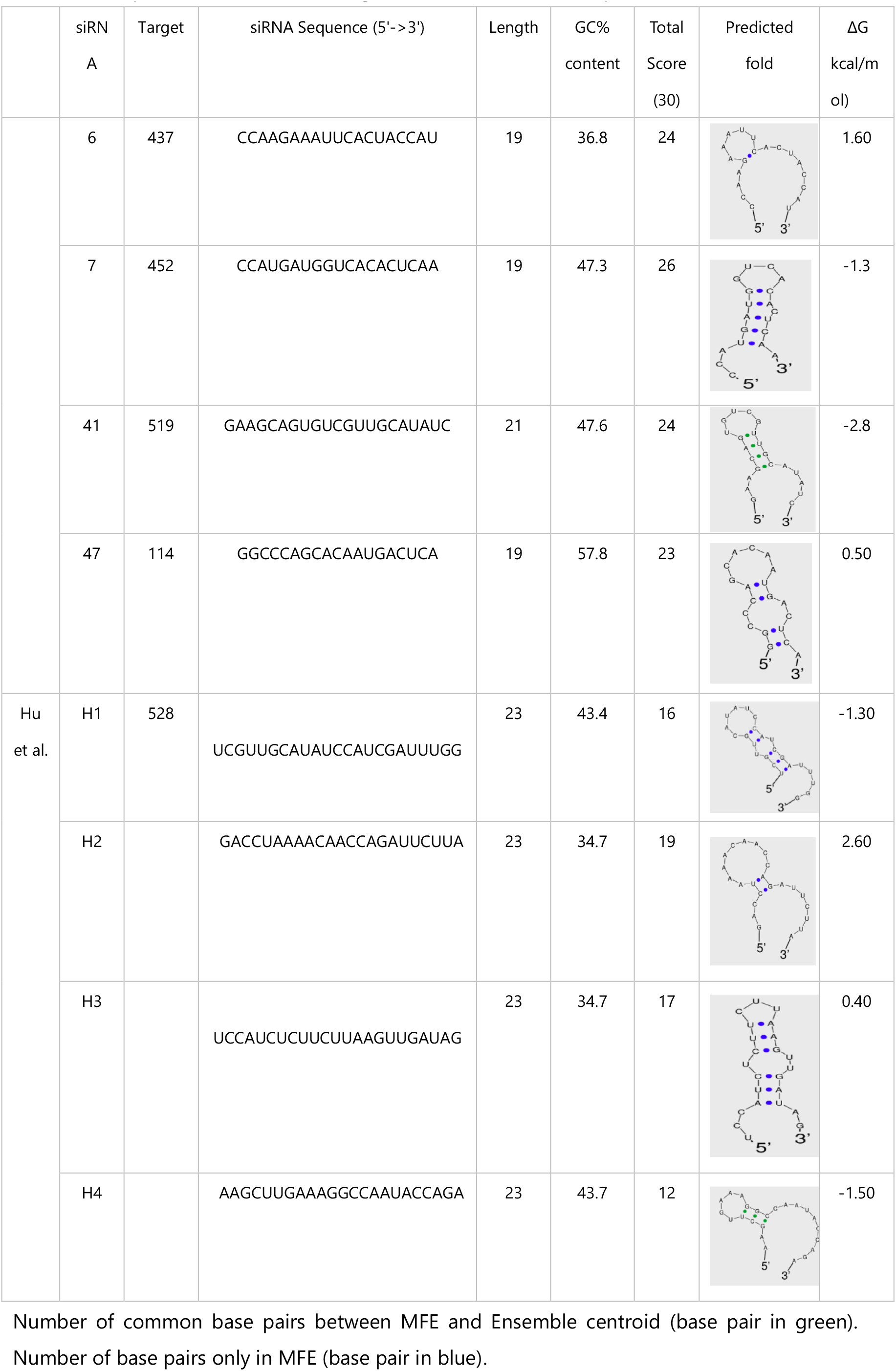
Comparison of the best scoring siRNA candidates with experimental candidates.

**Figure 2:**
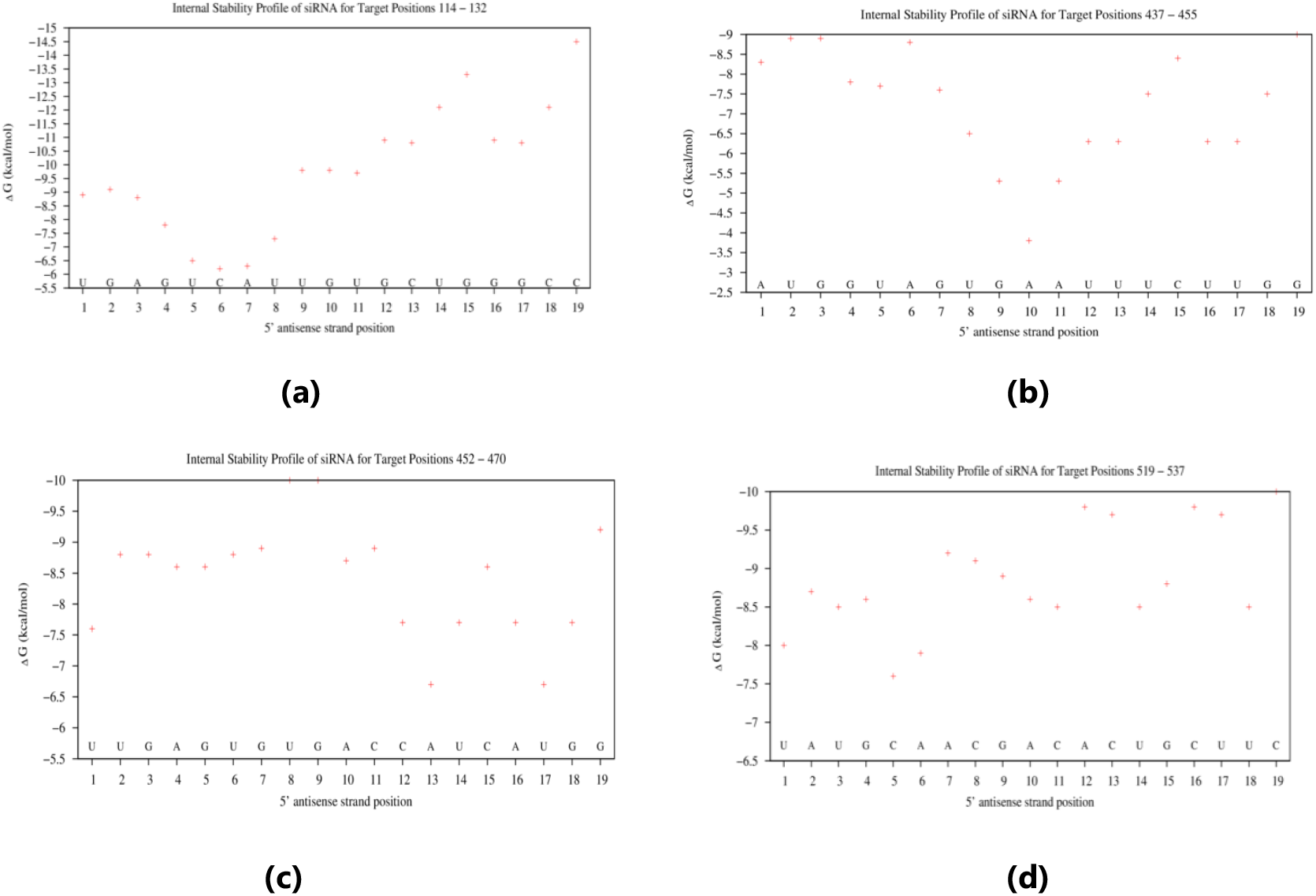
Internal stability profiles of siRNAs for particular target positions. The free energy of (a) siRNA-47 (b) sirna-6 (c) siRNA-7 and (d) siRNA-41 antisense strands for their target positions are shown.

Irrespective of how potential an siRNA is designed to be, gene silencing efficacy will be dependent on accessibility of the target site. As emphasized earlier, the stretch of nucleotides in the open reading frame for human gremlin-1 was used as the target for the siRNA design. Provided the first nucleotide begins the start codon, the target mRNA in this study had 552 nucleotides (Supplementary Fig. 1). Target binding site is another parameter that could define siRNA silencing efficacy [16]. Binding sites should occur at 50-100 nucleotides away from the start codon since regulatory proteins bind to those regions [31]. Further comprehensive analysis of the target site was performed with the Sirna algorithm of Sfold server. The module efficiently predicted the target site accessibility of the designed siRNA candidates. The probability of a target site available for siRNA binding was described with probability profiles (Figure 3). The folding patterns of the target sites were also described with loop specific probability profiles. In Figure 4, probabilities of the target sites exhibiting different folding patterns in the form of hairpin (Hplot), interior loops (Iplot) buge (Bplot) and multi-branched loop (Mplot) are shown.

**Figure 3:**
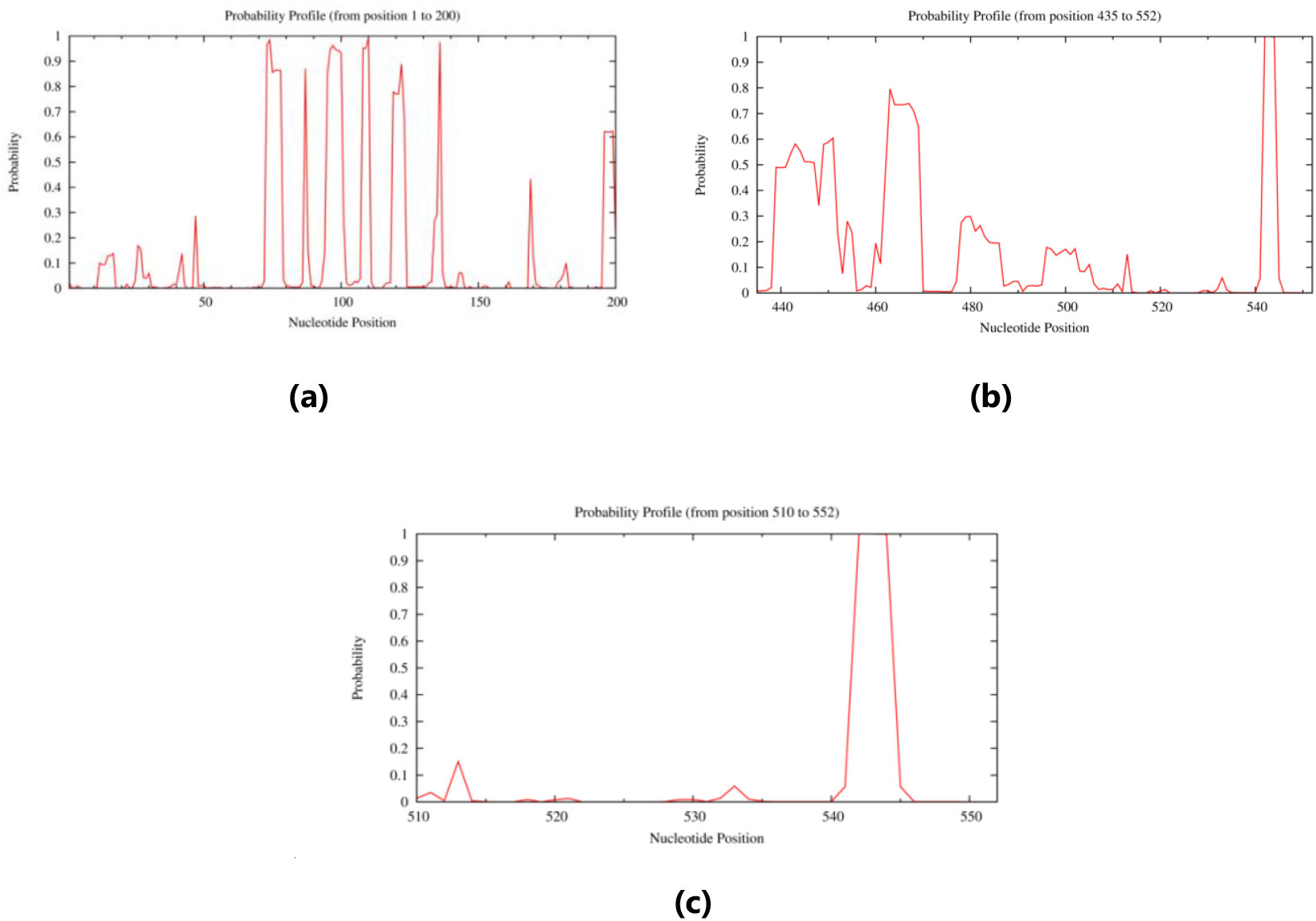
Probability profiles of target mRNA. The probability that a given stretch of nucleotides are all single-stranded and hence available for siRNA binding are shown for (a) target site 141 (siRNA-47); (b) target sites 437 (siRNA-6) and 452 (siRNA-7); and (c) target site 519 (siRNA-41).

Non-specific activities of siRNA in biological systems such as stimulation of immune responses and off-target effects limit efficiency of siRNA-mediated therapy. The occurrence of off-target effect is due to cross matching of siRNA to other mRNA transcripts apart from the target mRNA. A stringent basic local alignment search was performed to identify human genomic and transcript nucleotides that share homology with the designed siRNAs. Sequences that shared more than 75% similarity with the siRNA were considered for further evaluation to know whether the homology existed in the seed region. SiRNA-6, siRNA-7, and siRNA-47 all shared 100% similarity with human gremlin-1 transcript (Supplementary Fig. 3)

**Figure 4:**
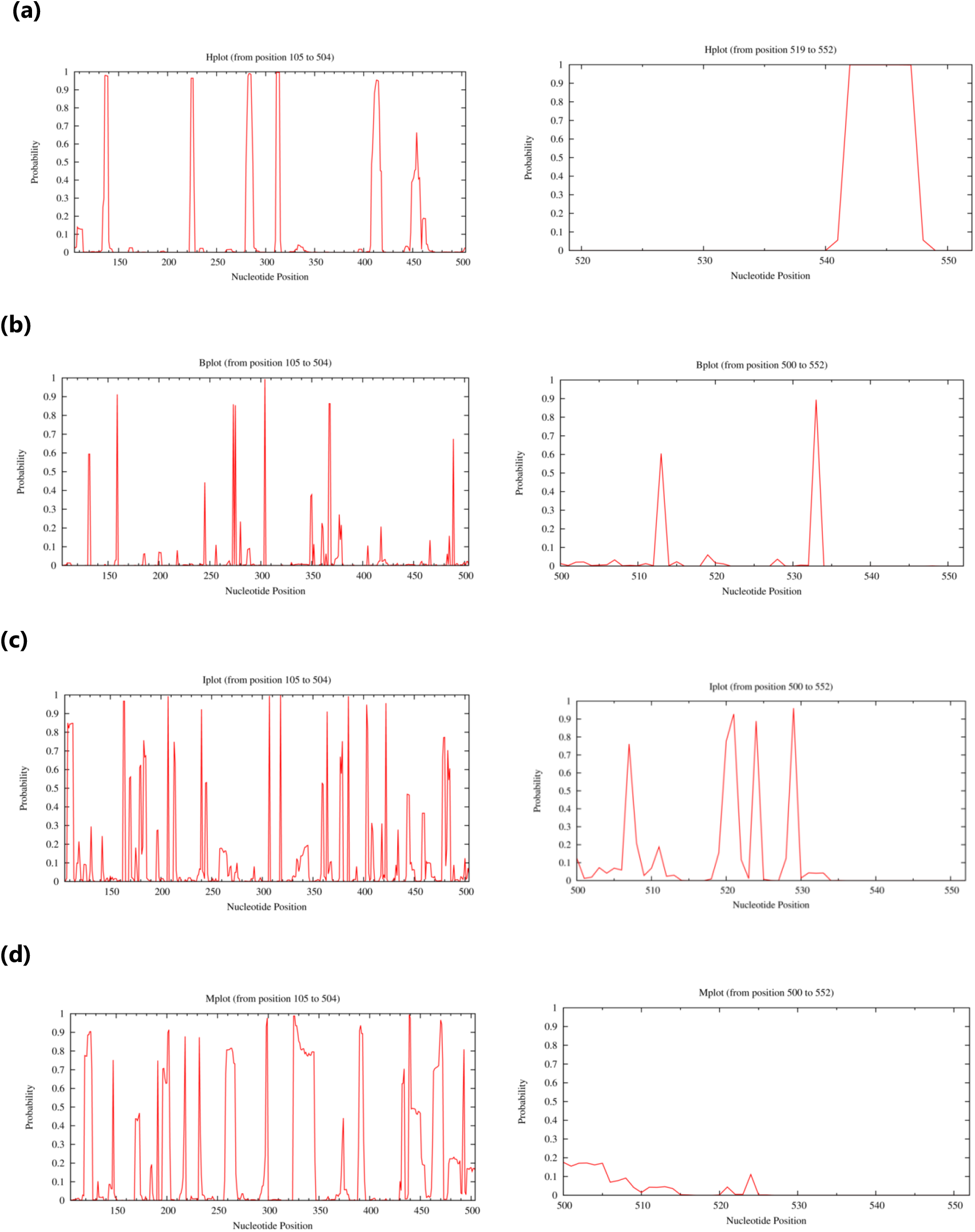
Loop specific profiles of target sites. (a) Displays the probability that a base is involved in hairpin loop (Hplot); (b) displays the probability that a base is in a bulge loop (Bplot); (c) displays the probability that a base is in an interior (internal) loop (Iplot); (d) displays the probability that a base is in a multibranched loop (Mplot).

## DISCUSSION

In this study, empirical guidelines for siRNA design and mRNA target accessibility have been comprehensively applied to select siRNA candidates that possess features for effective siRNA-mediated silencing of human gremlin-1. Gremlin-1 is a crucial signaling element for several biological pathways. During vertebrate limb development, gremlin-1 antagonizes bone morphogenetic proteins to maintain mechanisms for proper limb bud patterning [32]. Improper regulation and expression of gremlin-1 resulted in several defects and disorders [33, 34]. Gremlin-1 can induce angiogenic effects and fibrosis in organs, contributing to progression of cancer development of complex diseases [8]. The overexpression of gremlin-1 in cancer-associated fibroblasts has been reported [35].

RNAi-based therapy mediated by siRNA can offer a more specific inhibition of tumor growth than traditional chemotherapy [36] and has more potential than other methods of gene therapy [37]. Effective design of siRNA candidates is crucial, as non-specific siRNAs can produce unwanted effects. Several research groups reported sequence-based features of effective siRNAs and developed guidelines for siRNA design. In this study, guidelines proposed by three different research groups were used to select the best siRNA from a number of candidates designed using reliable computational algorithms. In Table 2, the highest scoring siRNA candidates are shown. These siRNA candidates achieved more than 75% based on the scoring criteria, hence, satisfy the sequence requirements for effective siRNA. Small interfering RNA molecules that satisfy the Ui-Tei guidelines were able to cause a significant reduction in firefly luciferase activity and also in chick embryo [22]. Meanwhile, siRNA candidates that obeyed the nucleotide composition demands of the Amarzguioui guidelines were described as functional [24]. Nucleotide sequence preference improve siRNA silencing efficiency as bases at particular positions define target recognition and cleavage [23]. For example, ‘A’ but not ‘G’ or ‘C’ is preferred at the 19^th^ position of the sense strand (Reynolds rule) in order to facilitate selection of siRNA entry into RISC [38]. Formation of stable intramoleular hairpin secondary structure may reduce availability of siRNA strands to mediate gene silencing. The secondary structures for all selected siRNA candidates were predicted with Sfold. The Srna algorithm of Sfold generate the secondary structure of the RNA with the minimum free energy (Table 2). However, [24] reported that intramolecular hairpin secondary structure of strand does not significantly affect siRNA efficacy.

Thermodynamics of siRNA duplex ends also define gene silencing potential [38]. To evaluate the free energy of the siRNA duplex ends, the Srna algorithm of Sfold server was used to compute the internal stabilities of siRNA for specific target sites (Figure 2). Physiologically, the dicer binds to the strand with less thermodynamic stability at the 5’ end, usually the antisense strand [22, 38]. According to [22], the free energy of the antisense strand of effective siRNA should not exceed −10kcal/mol. The free energy of antisense strands for the selected siRNAs in this study ranged from ≈-7.5kcal/mol to ≈-9kcal/mol. This agrees with reports that effective siRNAs are thermodynamically less stable at antisense 5’end.

The efficacy of the selected siRNA candidates ultimately depends on availability of their target sites. The probability that a target site is available for binding was computed with the Sirna module. The algorithm provides the probability profile for single-stranded regions in the target mRNA [28]. Target sites that are accessible have high probabilities of being single stranded. The probability profiles for all the selected siRNA target sites are shown in Figure 3. The loop specific probability profiles were generated to further evaluate the target site availability (Figure 4). The loop specific profiles represent the probability that a particular target site is not involved in hairpin, interior, bulge or multi-branched folds. The target sites for siRNA-6, siRNA-7 and siRNA-47 were predicted to be available. The target site for siRNA-47 (position 519-537) was not available for binding (Figure 1b). Moreover, the site was predicted to form an interior loop (Figure 4c).

To determine the specificity of the selected siRNA, local alignment search was performed against human genomic and transcript database (see Supplementary Fig. 3). This was to reduce off-target effects of siRNAs, a major challenge in RNAi-based therapy [16]. The three siRNA candidates shared a maximum of 78% similarity with other transcripts not recognized as gremlin-1. However, RNA transcripts are subjected to processing including splicing of intron segments to form mature mRNA. It is possible that the homologous nucleotides in the other transcripts are not components of an mRNA sequence. Furthermore, the mRNA target for this study was searched and aligned against human transcript and genome database and all the hits were identified as gremlin transcripts (see Supplementary Fig. 3d)

The selected siRNA candidates were compared to experimentally functional siRNAs, in order to evaluate the silencing potential of the designed siRNAs in this study. Reference [39] used gremlin-1 siRNAs to silence gremlin-1 mRNA in a study to determine the effect of gremlin-1 on BMP2-induced osteogenic differentiation of human bone marrow-derived mesenchymal stem cells. The siRNA sense strands used are designated as H1, H2, H3 and H4 in Table 2. The functional siRNAs (H1 and H2) were compared with the selected siRNAs in this study in terms of preferred sequence features for effective silencing. All three selected siRNAs in this study outscored H1 and H2, suggesting that they are effective siRNA candidates to silence human gremlin-1.

## CONCLUSION

Small interfering RNAs were designed to target human gremlin-1 using bioinformatics approach. Multiple computational algorithms were used to design potential siRNA candidates and the best candidates were selected based on empirical design rules and comprehensive target availability analysis. The selected candidates were then compared to experimentally functional siRNAs to suggest the efficacy of the designed siRNAs in this study. Out of 66 designed siRNA candidates, three were selected to be highly effective based on preferential sequence features for functional siRNAs, thermodynamics of siRNA duplexes and mRNA target availability. siRNA-6 (sense strand: 5’-CCAAGAAAUUCACUACCAU-3’), siRNA-7 (sense strand: 5’-CCAUGAUGGUCACACUCAA-3’) and siRNA-47 (sense strand: 5’-GGCCCAGCACAAUGACUCA) were predicted to be the most effective siRNA candidates to silence human gremlin-1 mRNA. Consideration for RNAi-based therapy is suggested because the siRNA candidates are predicted to exhibit reduced off-target effects after subjection to stringent similarity search against human genomics and transcript database.

