## Supplemental Tables and Figures for "Designing Effective small interfering RNA for Post-Transcriptional Silencing of Human GREM1: A Comprehensive Bioinformatics Approach"

**FIGURES AND TABLES**

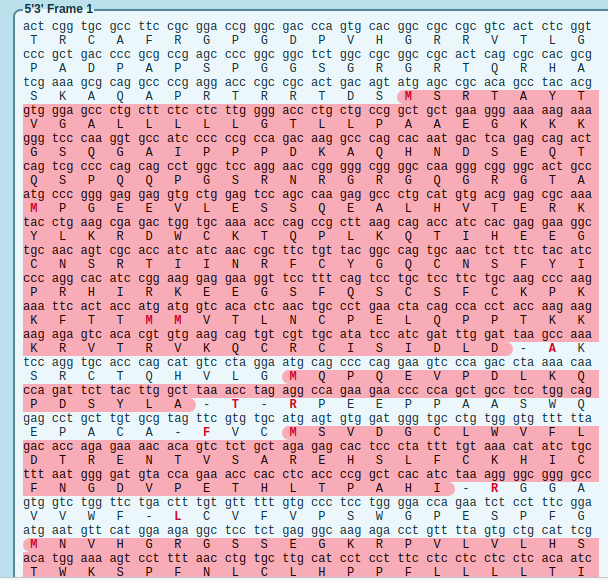

### Figure 1: Translation results of human gremlin-1 primary transcript (NM_001191323.2). Open reading frames are highlighted in red

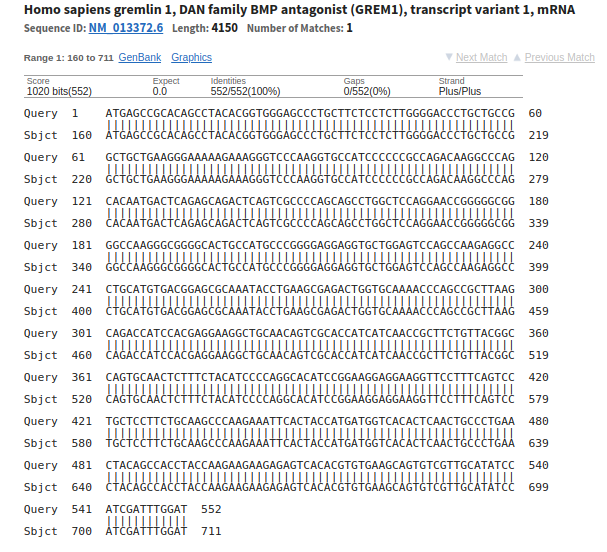

Figure 2: Alignment of mRNA target with gremlin-1 transcript 1 (NM_013372.7).

### Table 1: Predicted siRNA using different designing algorithms

| Computational tool | Target site start | siRNA sense (5'->3') |
| --- | --- | --- |
| BLOCK-iT™ RNAi Designer | 294 | GCTTAAGCAGACCATCCAC |
|  | 323 | GCAACAGTCGCACCATCAT |
|  | 425 | CCTTCTGCAAGCCCAAGAA |
|  | 428 | TCTGCAAGCCCAAGAAATT |
|  | 431 | GCAAGCCCAAGAAATTCAC |
|  | 437 | CCAAGAAATTCACTACCAT |
|  | 452 | CCATGATGGTCACACTCAA |
|  | 475 | CCTGAACTACAGCCACCTA |
|  | 486 | GCCACCTACCAAGAAGAAG |
|  | 528 | TCGTTGCATATCCATCGAT |
| OligoWalk | 326 | UUGAUGAUGGUGCGACUGU |
|  | 487 | UCUUCUUCUUGGUAGGUGG |
|  | 359 | UAGAAAGAGUUGCACUGGC |
|  | 61 | UCUUUUUCCCUUCAGCAGC |
|  | 493 | UGACUCUCUUCUUCUUGGU |
|  | 465 | UAGUUCAGGGCAGUUGAGU |
|  | 440 | AUCAUGGUAGUGAAUUUCU |
|  | 425 | UUCUUGGGCUUGCAGAAGG |
|  | 62 | UUCUUUUUCCCUUCAGCAG |
|  | 485 | UUCUUCUUGGUAGGUGGCU |
|  | 530 | AAAUCGAUGGAUAUGCAAC |
|  | 514 | AACGACACUGCUUCACACG |
|  | 59 | UUUUUCCCUUCAGCAGCCG |
|  | 362 | AUGUAGAAAGAGUUGCACU |
|  | 426 | UUUCUUGGGCUUGCAGAAG |
|  | 338 | UAACAGAAGCGGUUGAUGA |
|  | 491 | ACUCUCUUCUUCUUGGUAG |
|  | 63 | UUUCUUUUUCCCUUCAGCA |
|  | 529 | AAUCGAUGGAUAUGCAACG |
|  | 490 | CUCUCUUCUUCUUGGUAGG |
|  | 244 | AUUUGCGCUCCGUCACAUG |
|  | 534 | AUCCAAAUCGAUGGAUAUG |
|  | 524 | AUGGAUAUGCAACGACACU |
|  | 518 | AUGCAACGACACUGCUUCA |
|  | 396 | AAAGGAACCUUCCUCCUUC |
|  | 123 | AGUCUGCUCUGAGUCAUUG |
|  | 108 | AUUGUGCUGGGCCUUGUCU |
| siRNA Wizard | 334 | ACCATCATCAACCGCTTCTGT |
|  | 358 | GGCCAGTGCAACTCTTTCTAC |
|  | 463 | ACACTCAACTGCCCTGAACTA |
|  | 519 | GAAGCAGTGTCGTTGCATATC |
| i-Score Designer | 452 | CCAUGAUGGUCACACUCAA |
|  | 519 | GAAGCAGUGUCGUUGCAUA |
|  | 61 | GCUGCUGAAGGGAAAAAGA |
|  | 216 | GGUGCUGGAGUCCAGCCAA |
|  | 63 | UGCUGAAGGGAAAAAGAAA |
|  | 114 | GGCCCAGCACAAUGACUCA |
|  | 57 | GCCGGCUGCUGAAGGGAAA |
|  | 422 | GCUCCUUCUGCAAGCCCAA |
|  | 475 | CCUGAACUACAGCCACCUA |
|  | 243 | GCAUGUGACGGAGCGCAAA |
| RNAxs | 377 | UAGAAAGAGUUGCACUGGC |
|  | 503 | UUCUUCUUGGUAGGUGGCU |
|  | 504 | CUUCUUCUUGGUAGGUGGC |
|  | 446 | AAUUUCUUGGGCUUGCAGA |
|  | 455 | AUGGUAGUGAAUUUCUUGG |
|  | 493 | UAGGUGGCUGUAGUUCAGG |
|  | 454 | UGGUAGUGAAUUUCUUGGG |
|  | 492 | AGGUGGCUGUAGUUCAGGG |
|  | 375 | GAAAGAGUUGCACUGGCCG |
|  | 125 | UUGUGCUGGGCCUUGUCUG |
|  | 299 | UUAAGCGGCUGGGUUUUGC |
|  | 374 | AAAGAGUUGCACUGGCCGU |
|  | 356 | UAACAGAAGCGGUUGAUGA |
|  | 376 | AGAAAGAGUUGCACUGGCC |
|  | 505 | UCUUCUUCUUGGUAGGUGG |

### Table 2: Scoring of siRNA using proposed design guidelines.

|  | | | | Scoring | | | |
| --- | --- | --- | --- | --- | --- | --- | --- |
| siRNA | Target site start | siRNA Sequence (5'->3') | Length | Ui-Tei (8) | Amarzguioui (12) | Reynolds (10) | Total (30) |
| 1 | 294 | GCUUAAGCAGACCAUCCAC | 19 | 6 | 8 | 4 | 18 |
| 2 | 323 | GCAACAGUCGCACCAUCAU | 19 | 5 | 8 | 6 | 19 |
| 3 | 425 | CCUUCUGCAAGCCCAAGAA | 19 | 6 | 6 | 6 | 18 |
| 4 | 428 | UCUGCAAGCCCAAGAAAUU | 19 | 5 | 6 | 6 | 17 |
| 5 | 431 | GCAAGCCCAAGAAAUUCAC | 19 | 5 | 6 | 6 | 17 |
| **6** | **437** | **CCAAGAAAUUCACUACCAU** | **19** | **6** | **10** | **8** | **24** |
| **7** | **452** | **CCAUGAUGGUCACACUCAA** | **19** | **8** | **8** | **10** | **26** |
| 9 | 486 | GCCACCUACCAAGAAGAAG | 19 | 4 | 4 | 4 | 12 |
| 10 | 528 | UCGUUGCAUAUCCAUCGAU | 19 | 4 | 6 | 6 | 16 |
| 11 | 326 | UUGAUGAUGGUGCGACUGU | 19 | 3 | 6 | 6 | 15 |
| 12 | 487 | UCUUCUUCUUGGUAGGUGG | 19 | 1 | 4 | 6 | 11 |
| 13 | 359 | UAGAAAGAGUUGCACUGGC | 19 | 3 | 4 | 6 | 13 |
| 14 | 61 | UCUUUUUCCCUUCAGCAGC | 19 | 0 | 4 | 2 | 6 |
| 15 | 493 | UGACUCUCUUCUUCUUGGU | 19 | 6 | 6 | 8 | 20 |
| 16 | 465 | UAGUUCAGGGCAGUUGAGU | 19 | 5 | 4 | 4 | 13 |
| 17 | 440 | AUCAUGGUAGUGAAUUUCU | 19 | 7 | 4 | 6 | 17 |
| 18 | 425 | UUCUUGGGCUUGCAGAAGG | 19 | 1 | 2 | 6 | 9 |
| 19 | 62 | UUCUUUUUCCCUUCAGCAG | 19 | 0 | 4 | 2 | 6 |
| 20 | 485 | UUCUUCUUGGUAGGUGGCU | 19 | 1 | 4 | 2 | 7 |
| 21 | 530 | AAAUCGAUGGAUAUGCAAC | 19 | 4 | 4 | 4 | 12 |
| 22 | 514 | AACGACACUGCUUCACACG | 19 | 0 | 6 | 4 | 10 |
| 23 | 59 | UUUUUCCCUUCAGCAGCCG | 19 | 1 | 2 | 4 | 7 |
| 24 | 362 | AUGUAGAAAGAGUUGCACU | 19 | 5 | 8 | 4 | 17 |
| 25 | 426 | UUUCUUGGGCUUGCAGAAG | 19 | 0 | 2 | 4 | 6 |
| 26 | 338 | UAACAGAAGCGGUUGAUGA | 19 | 5 | 6 | 8 | 19 |
| 27 | 491 | ACUCUCUUCUUCUUGGUAG | 19 | 5 | 4 | 6 | 15 |
| 28 | 63 | UUUCUUUUUCCCUUCAGCA | 19 | 3 | 6 | 4 | 13 |
| 29 | 529 | AAUCGAUGGAUAUGCAACG | 19 | 3 | 6 | 4 | 13 |
| 30 | 490 | CUCUCUUCUUCUUGGUAGG | 19 | 3 | 6 | 6 | 15 |
| 31 | 244 | AUUUGCGCUCCGUCACAUG | 19 | 2 | 6 | 6 | 14 |
| 32 | 534 | AUCCAAAUCGAUGGAUAUG | 19 | 4 | 4 | 4 | 12 |
| 33 | 524 | AUGGAUAUGCAACGACACU | 19 | 5 | 6 | 6 | 17 |
| 34 | 518 | AUGCAACGACACUGCUUCA | 19 | 5 | 8 | 6 | 19 |
| 35 | 396 | AAAGGAACCUUCCUCCUUC | 19 | 3 | 6 | 6 | 15 |
| 36 | 123 | AGUCUGCUCUGAGUCAUUG | 19 | 3 | 2 | 6 | 11 |
| 37 | 108 | AUUGUGCUGGGCCUUGUCU | 19 | 3 | 4 | 6 | 13 |
| 38 | 334 | ACCAUCAUCAACCGCUUCUGU | 21 | 4 | 4 | 6 | 14 |
| 39 | 358 | GGCCAGUGCAACUCUUUCUAC | 21 | 6 | 8 | 6 | 20 |
| 40 | 463 | ACACUCAACUGCCCUGAACUA | 21 | 3 | 2 | 10 | 15 |
| **41** | **519** | **GAAGCAGUGUCGUUGCAUAUC** | **21** | **6** | **10** | **8** | **24** |
| 44 | 61 | GCUGCUGAAGGGAAAAAGA | 19 | 5 | 6 | 6 | 17 |
| 45 | 216 | GGUGCUGGAGUCCAGCCAA | 19 | 3 | 8 | 4 | 15 |
| 46 | 63 | UGCUGAAGGGAAAAAGAAA | 19 | 5 | 6 | 6 | 17 |
| **47** | **114** | **GGCCCAGCACAAUGACUCA** | **19** | **5** | **12** | **6** | **23** |
| 48 | 57 | GCCGGCUGCUGAAGGGAAA | 19 | 6 | 6 | 8 | 20 |
| 49 | 422 | GCUCCUUCUGCAAGCCCAA | 19 | 3 | 8 | 4 | 15 |
| 50 | 475 | CCUGAACUACAGCCACCUA | 19 | 3 | 10 | 6 | 19 |
| 51 | 243 | GCAUGUGACGGAGCGCAAA | 19 | 3 | 8 | 6 | 17 |
| 52 | 377 | UAGAAAGAGUUGCACUGGC | 19 | 3 | 4 | 6 | 13 |
| 53 | 503 | UUCUUCUUGGUAGGUGGCU | 19 | 1 | 4 | 2 | 7 |
| 54 | 504 | CUUCUUCUUGGUAGGUGGC | 19 | 2 | 4 | 4 | 8 |
| 55 | 446 | AAUUUCUUGGGCUUGCAGA | 19 | 5 | 8 | 4 | 17 |
| 56 | 455 | AUGGUAGUGAAUUUCUUGG | 19 | 5 | 6 | 4 | 15 |
| 57 | 493 | UAGGUGGCUGUAGUUCAGG | 19 | 2 | 4 | 4 | 10 |
| 58 | 454 | UGGUAGUGAAUUUCUUGGG | 19 | 3 | 4 | 4 | 11 |
| 59 | 492 | AGGUGGCUGUAGUUCAGGG | 19 | 1 | 4 | 6 | 11 |
| 60 | 375 | GAAAGAGUUGCACUGGCCG | 19 | 2 | 6 | 4 | 12 |
| 61 | 125 | UUGUGCUGGGCCUUGUCUG | 19 | 2 | 4 | 4 | 10 |
| 62 | 299 | UUAAGCGGCUGGGUUUUGC | 19 | 5 | 2 | 8 | 15 |
| 63 | 374 | AAAGAGUUGCACUGGCCGU | 19 | 3 | 8 | 4 | 15 |
| 64 | 356 | UAACAGAAGCGGUUGAUGA | 19 | 5 | 6 | 8 | 19 |
| 65 | 376 | AGAAAGAGUUGCACUGGCC | 19 | 1 | 2 | 6 | 9 |
| 66 | 505 | UCUUCUUCUUGGUAGGUGG | 19 | 1 | 4 | 6 | 11 |

siRNAs in bold are selected high scoring candidates.

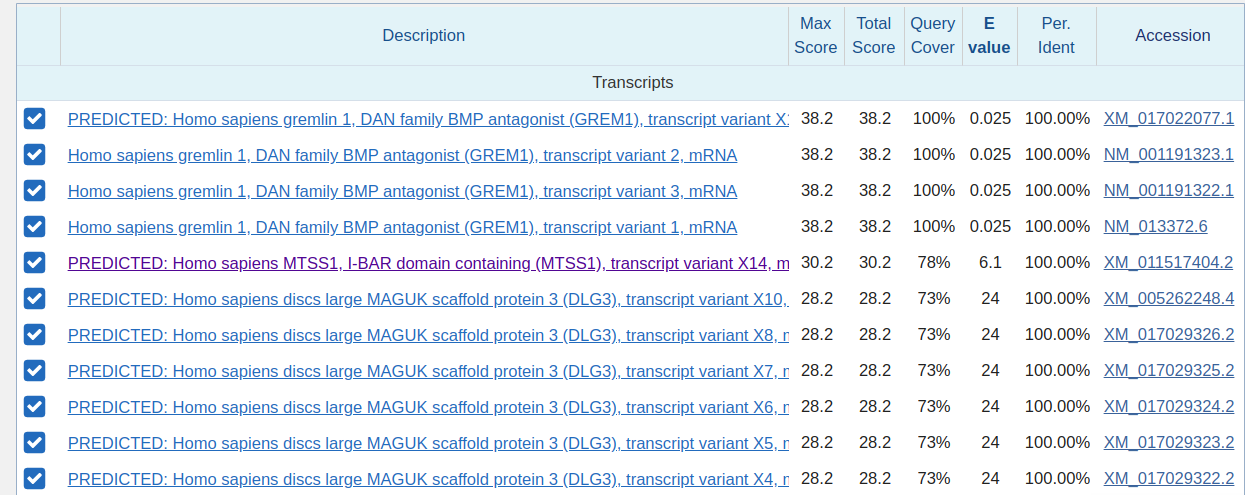

**(a)**

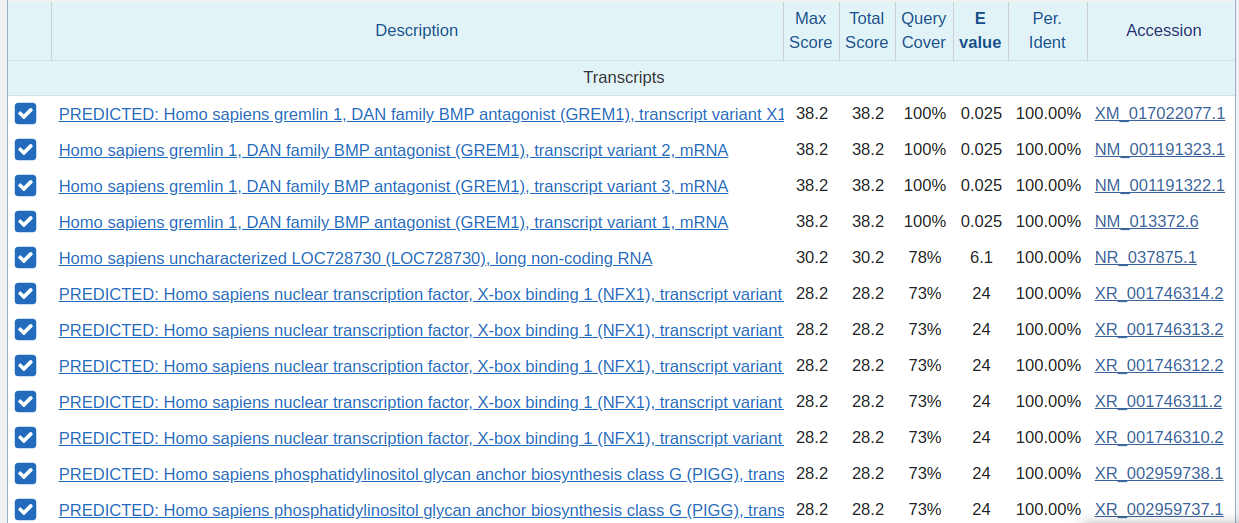

**(b)**

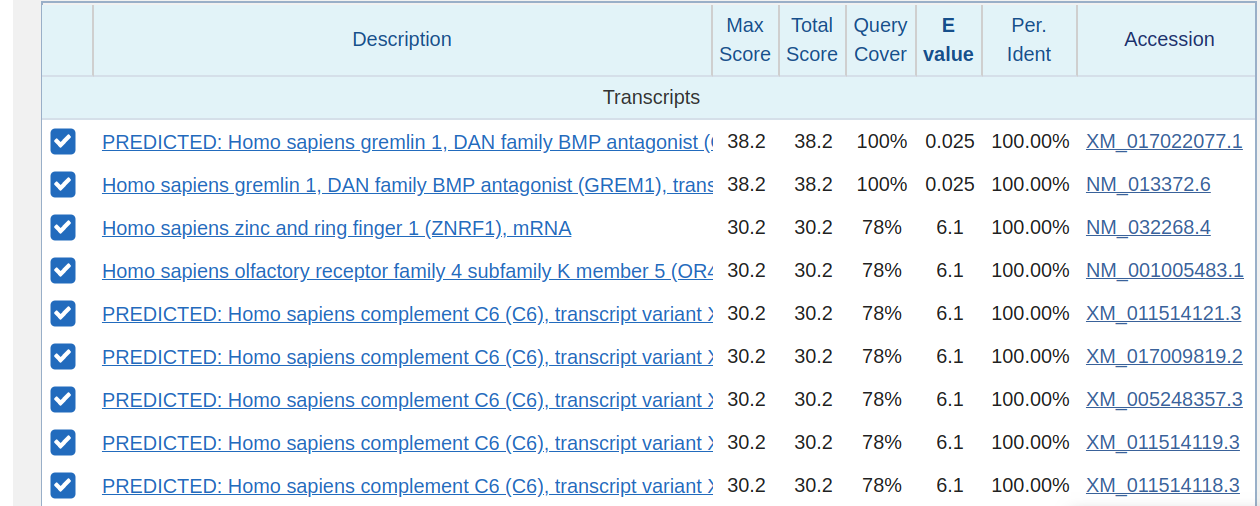

**(c)**

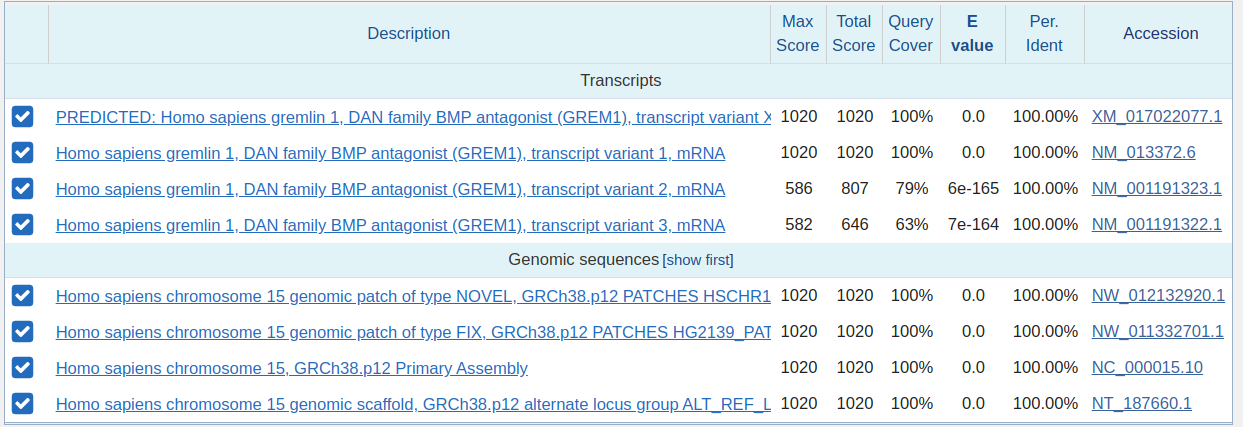

**(d)**

### Figure 3: Blast hits for nucleotide similarity seraches. (a) displays blast hits for siRNA-6; (b) displays blast hits for siRNA-7; (c) displays blast hits for siRNA-47; (d) displays blast hits for mRNA target.
